# Song complexity in relation to repertoire size and phonological syntax in the breeding song of Purple Sunbird

**DOI:** 10.1101/2021.03.20.436261

**Authors:** Sonam Chorol, Manjari Jain

## Abstract

There are multiple measures for bird song complexity such as repertoire size, phonological or compositional syntax and complex vocal mechanism (CVM). We examined these in an old-world passerine, Purple Sunbird. First, we identified and acoustically characterised the repertoire size (of notes and phrases). We then assessed positional fidelity and ordering of notes within phrases. We found 23 distinct notes by aural-visual inspection of spectrograms which was validated using a Classification and Regression Tree based on 5 acoustic parameters. These notes combined in different iterations to form 30 different phrases. Phrases comprised of an overall structure with an introductory note (prefix) at the onset, followed by notes occurring at specific positions within the phrase body, and terminated with a trill (suffix syllable(s)). Prefix was present in 93% of phrases whereas suffix syllable(s) occurred in 27% of phrases only. We found that notes exhibited positional fidelity and combined in specific order to form a phrase. This is indicative of underlying phonological syntax that limits the ways in which notes combine to form phrase and offers insights to song complexity. Finally, we found that suffix syllables exhibit the presence of mini-breath (very short inter-note interval) which are known to be produced by CVM.

## Introduction

The total set of vocalizations that a species possesses and uses in different behavioural contexts is regarded as the vocal repertoire of a species (Searcy 1992). A large vocal repertoire depicts higher complexity in communication (Blumstein and Armitage 1997; McComb and Semple 2005). This phenomenon has been studied extensively in primates as well as in avian species (Range and Fischer 2004; Gustison et al. 2012). Structural complexity of acoustic signals exists not only in the variety of vocalizations but also in the manner in which vocalizations are organised and composed (Hailman and Ficken 1986). In avian vocalizations the smallest acoustic unit is referred to as a ‘note’ or ‘element’ which can combine, sometimes following different ordering rules, to form higher order vocal units called ‘phrase’. Finally, songs can be composed of repetition of a single element (note) or phrases (the same or different kind).

A phrase may be composed of a single note repeated multiple times or different notes occurring in a defined sequence. Moreover, notes may be shared between two or more phrases (Kroodsma 1977) and construction of phrases may follow certain patterns. The combinatorial rule that governs the construction of the signal from subset of signals is termed as syntax and it could be compositional or phonological syntax. The rules for arranging smaller meaningful units into a higher meaningful signal is called compositional syntax (Berwick et al. 2011). Altering this sequence may change the meaning of the signal (Berwick et al. 2011). The presence of compositional syntax is well known in human language where meaningful signalling units ‘words’ combines to form a higher meaningful signal ‘phrase’. In avian systems, the presence of the same is reported where ‘alert call’ and ‘recruitment call’ combine to form ‘mobbing call’ (Suzuki et al. 2016; Engesser et al. 2016). On the other hand, bird song is a combination of meaningless units (notes) into phrases which too do not have a defined meaning. However, the notes may combine in a defined and non-random manner to build the phrase. Thus, the construction of song phrases may follow a combinatorial rule thereby exhibits a phonological syntax (Berwick et al. 2011). Moreover, bird song comprises of phrases and each phrase may be initiated with an introductory note(s) - a stereotypic note (s) which is produced at the onset of the song phrase, followed by series of notes, structurally similar or distinct, which form the main component of a phrase (phrase body) (Williams 2004; Roach et al. 2012; Nelson and Soha 2004). A song can end with a terminal trill that includes a series of rapidly produced notes (Nelson and Soha 2004). The ordering of notes within a phrase following a non-random positional occurrence is indicative of phonological syntax and is another aspect of structural complexity.

Furthermore, song complexity also depends upon different vocal mechanisms. For instance, some birds can produce complex vocalizations with two temporally overlapping notes of different frequencies. These complex vocalizations are produced using both sides of the syrinx by expiring half air from one side of the syrinx at a certain frequency and the other half from the other side at a different frequency (Suthers 2004). These notes are separated by a short time interval (< 20 ms) known as a ‘mini-breath’. The production of such a syllable (acoustic unit composed of 2-3 notes) is expected to require precise bilateral motor coordination within the syrinx and between various muscles (Suthers 2004; Suthers et al. 2012).

In this study, we examined song complexity in an Old-World passerine, Purple Sunbird (*Cinnyris asiaticus*). It belongs to the family Necteriniidae and occurs in West Asia, throughout the Indian subcontinent and into Southeast Asia (Ali and Ripley 1983). Purple Sunbirds are sexually dimorphic. Females are “olive brown dorsally with a yellowish underside whereas eclipse (nonbreeding) males have a distinct median line down the centre of throat and breast” (Ali and Ripley 1983). Males gain bright, metallic blue-green coloured plumage during the breeding season which ranges from April-June in Northern India (Ali and Ripley 1983). During breeding season, male sings multiple song with ringing metallic notes and increases the loudness of song under noisy conditions (Singh et al. 2019). In this study, we aimed to examine the breeding song complexity of Purple Sunbird at multiple levels. The objectives were as follows: a) to determine the repertoire size in terms of notes and phrases that compose the breeding song b) to examine the presence of phonological syntax with respect to combinatorial rules underlying phrase structuring and composition and c) to investigate evidence for complex vocal mechanism (CVM) indicated by presence of ‘sexy syllables’ in the vocal repertoire.

## Methods

### Sampling location

Sampling was done during the breeding season between May-August 2017 in IISER Mohali campus (30.6650° N, 76.7300° E), Punjab. The locality is a semi-urban, subtropical region which falls under ‘Cwa’ category (climate that is variable throughout the year with a hot summer and cold, dry winter separated by a brief period of tropical monsoon) of Koppen-Geiger climate classification (Kottek et al. 2006). The vegetation is predominantly grassy with intermittent canopy of dry deciduous mixed with evergreen trees including host plants such as *Callistemon linearis, Delonix regia, Habiscus* sp., *Plumeria* sp., *Lantena camara, Cascabela thevetia, Hamelia petens*, etc.

### Song recording and analyses

A total of 3026 notes and 241phrases were analysed from the songs of Purple Sunbird recorded over four months. Recordings were made opportunistically using a solid-state recorder (Marantz PMD661-MKII; frequency response: 20 Hz – 20 kHz), connected to a super-cardioid shotgun microphone (Sennheiser ME66 with K6 PM; frequency response: 40 Hz to 20 kHz), covered with a foam windscreen (Sennheiser MZW66). All vocalizations were recorded at a sampling rate of 44.1 kHz with 16-bit accuracy. All recordings were processed in Raven Pro 1.5 (Cornel Lab of Ornithology) and spectrogram generated using Hann-window with 512 window size and 50% overlap. Songs were categorised preliminarily based on aural-visual inspection and a catalogue of all note types was generated and notes were annotated as A, B, C.... W. These were then subjected to detailed acoustic analyses based on 1 temporal (note duration) and 4 spectral (frequency 5%, frequency 95%, frequency bandwidth 90% and peak frequency) parameters. Note duration was calculated as the time duration between onset and offset of a note. Frequency 5% and 95% represent frequencies that lie at 5% and 95% of the energy of a sound signal respectively and bandwidth 90% was the difference between frequency 5% and 95%. Peak frequency represents the frequency with maximum energy. Analyses of phrases was carried out by determining the note type and their respective position within the note. Phrases with similar note composition and ordering but with variation in repetition of a note type were considered as the same. This gives a conservative estimate of the repertoire size at the note and phrase level.

### Validation of classical analyses with CART

To validate the repertoire size calculated based on aural-visual inspection method, note classification was performed using ‘Classification and Regression Tree’ (CART; ‘rpart’ package; Therneau et al. 2018) in R 4.0.3 (R development core team 2008) following the protocol of Garland et al. 2015. We reduced the sample size to an upper limit of N=50 per note to eliminate the over-representation of notes with large sample sizes by random sampling. The data was partitioned at 3:1 as training and test data set. The splitting of nodes was based on the ‘Gini index’ which reduce the impurities at terminal nodes as it accounts for probability of misclassification. It may be noted that analysis performed using CART is robust to outliers, non-normal and non-independent data (Breiman et al. 1984).

### Evidence of phonological syntax

A total of 241 phrases were analysed to determine the phrase structure (presence and absence of introductory note (prefix) and terminal trill (suffix)). Moreover, underlying combinatorial rules that dictate note occurrence and ordering of notes would be indicative of phonological syntax. Towards this, we examined positional fidelity of notes by calculating the frequency of occurrence of each note on a specific position in phrase. We then plotted a heat matrix (using r package heatmaply; Galili et al. 2017) that depicts the probability of occurrence of every note in every possible position within a phrase. This was then compared to a null matrix to examine if the ‘observed’ probability of occurrence of notes in specific positions is non-random. Towards this we generated 100 matrices where the position of every note was randomly assigned within a phrase. The averaged value of this was then used to generate a null matrix, based on which an ‘expected’ heat plot was generated. The ‘observed’ heat-plot was then compared visually with the ‘expected’ plot to verify positional fidelity.

### Evidence of complex vocal mechanism

The temporal arrangement of notes in phrases was analysed based on the inter-note time interval. The inter-note interval is the duration between the offset of one note to the onset of the subsequent note within a phrase. We analysed inter-note interval position up to 20 positions as sample size of phrase comprising of >21 notes were less. These positions were marked in ascending order. Inter-note interval between prefix and the first note of body was always marked as 1. In phrases where prefix was absent, marking of inter-note time interval started from 2. A total of 2654 inter-note time intervals were analysed. To examine the differences in inter-note interval between different components of a phrase, we categorised inter-note interval into 3 groups - between prefix and body (PB), within body (B) and within suffix syllable (SS). This was done for all 241 phrases and the average values for the body and suffix region (since there would be >3 inter-note interval values in these regions) for a given phrase was taken for further analyses.

### Statistical Analysis

Statistical tests were performed in R 4.0.3 (R development core team 2008). Differences in frequency of occurrence of prefix and suffix syllable in the phrase were tested by χ^2^ test (using chisq.test function of the r package). We examined the correlation between inter-note interval and interval position of phrase checked by Pearson’s correlation (using cor.test function of the r package). Generalised Linear Model (GLM) fitted with Poisson as a family function (*glm* function of the r package) was used to compare inter-note interval between each note in a phrase. Post-hoc comparison of inter-note interval between 3 categories (PB, B and SS) was done by Mann-Whitney U (MW U) test (using wilcox.test function of the r package).

## Results

### Note classification and song repertoire

Aural-visual analyses of notes resulted in the identification of 23 different notes in the songs of Purple Sunbird (Table 1). This result was upheld by the Classification and Regression Tree (CART) analyses which classified notes into 23 terminal nodes with an accuracy of 78.08%. The first branch of the tree was based on delta time (note duration), which separated 5 smallest notes from the rest of the note types. At each branching the acoustic parameter responsible for the branching was identified and a total of 22 distinct notes were found (Figure 1).

**Figure 1.**
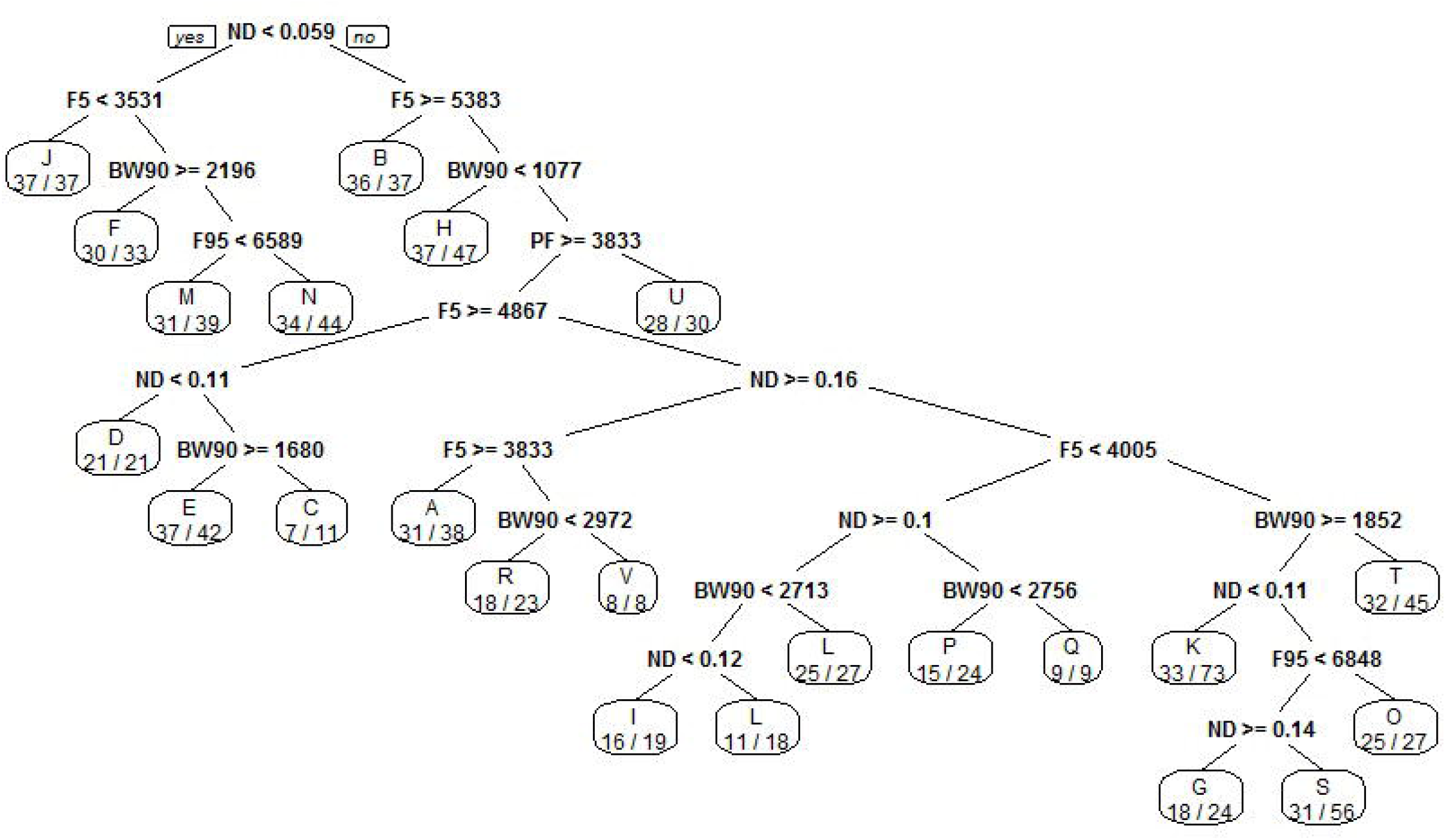
Classification of notes obtained from CART. The variables used at each split in the tree are listed, along with the criteria (<, >, or =). ND = note duration, F5 = frequency 5%, F95 = frequency 95%, BW90 = bandwidth 90% and PF = peak frequency. Right side of split agrees the criteria of splitting variable and left side does not agree. The terminal node represents the final classification of tree along with proportion of correctly classified values.

**Table 1.** Mean ± SD of 5 acoustic parameters for 23 notes obtained in the vocalization of Purple sunbird. N corresponds to sample size.

| S.No | Note ID | N | Note duration (s) | Frequency 5% (kHz) | Frequency 95% (kHz) | Bandwidth 90% (kHz) | Peak frequency (kHz) |
| --- | --- | --- | --- | --- | --- | --- | --- |
| 1 | A | 249 | $0.17 \pm 0.01$ | $4.00 \pm 0.20$ | $5.68 \pm 0.26$ | $1.68 \pm 0.37$ | $4.98 \pm 0.36$ |
| 2 | B | 64 | $0.15 \pm 0.02$ | $5.65 \pm 0.23$ | $7.05 \pm 0.14$ | $1.40 \pm 0.24$ | $6.90 \pm 0.17$ |
| 3 | C | 24 | $0.19 \pm 0.01$ | $4.96 \pm 0.10$ | $6.04 \pm 0.15$ | $1.08 \pm 0.15$ | $5.65 \pm 0.21$ |
| 4 | D | 267 | $0.07 \pm 0.01$ | $4.98 \pm 0.24$ | $6.67 \pm 0.31$ | $1.69 \pm 0.38$ | $6.12 \pm 0.42$ |
| 5 | E | 165 | $0.14 \pm 0.01$ | $5.02 \pm 0.07$ | $6.91 \pm 0.08$ | $1.89 \pm 0.09$ | $5.63 \pm 0.74$ |
| 6 | F | 116 | $0.05 \pm 0.01$ | $4.51 \pm 0.48$ | $7.03 \pm 0.38$ | $2.52 \pm 0.41$ | $6.47 \pm 0.74$ |
| 7 | G | 90 | $0.15 \pm 0.01$ | $4.33 \pm 0.21$ | $6.49 \pm 0.14$ | $2.16 \pm 0.29$ | $4.91 \pm 0.55$ |
| 8 | H | 155 | $0.15 \pm 0.05$ | $4.86 \pm 0.31$ | $5.70 \pm 0.32$ | $0.84 \pm 0.29$ | $5.32 \pm 0.32$ |
| 9 | I | 56 | $0.11 \pm 0.01$ | $4.11 \pm 0.24$ | $6.48 \pm 0.27$ | $2.37 \pm 0.22$ | $5.56 \pm 0.68$ |
| 10 | J | 115 | $0.04 \pm 0.01$ | $3.22 \pm 0.18$ | $4.72 \pm 0.41$ | $1.49 \pm 0.44$ | $3.81 \pm 0.36$ |
| 11 | K | 223 | $0.10 \pm 0.01$ | $4.37 \pm 0.20$ | $6.49 \pm 0.17$ | $2.58 \pm 0.24$ | $5.78 \pm 0.96$ |
| 12 | L | 720 | $0.12 \pm 0.01$ | $3.69 \pm 0.24$ | $6.39 \pm 0.36$ | $2.70 \pm 0.40$ | $4.86 \pm 0.63$ |
| 13 | M | 249 | $0.05 \pm 0.01$ | $4.97 \pm 0.23$ | $6.10 \pm 0.37$ | $1.13 \pm 0.41$ | $5.62 \pm 0.28$ |
| 14 | N | 13 | $0.04 \pm 0.00$ | $5.51 \pm 0.23$ | $7.20 \pm 0.14$ | $1.69 \pm 0.31$ | $6.63 \pm 0.23$ |
| 15 | O | 213 | $0.14 \pm 0.01$ | $4.54 \pm 0.18$ | $6.85 \pm 0.34$ | $2.31 \pm 0.35$ | $5.29 \pm 0.40$ |
| 16 | P | 33 | $0.07 \pm 0.01$ | $3.87 \pm 0.19$ | $5.88 \pm 0.31$ | $2.02 \pm 0.41$ | $4.83 \pm 0.45$ |
| 17 | Q | 16 | $0.09 \pm 0.01$ | $3.93 \pm 0.15$ | $6.91 \pm 0.16$ | $2.98 \pm 0.17$ | $6.15 \pm 0.53$ |
| 18 | R | 6 | $0.17 \pm 0.01$ | $3.70 \pm 0.09$ | $6.23 \pm 0.20$ | $2.53 \pm 0.26$ | $4.62 \pm 0.35$ |
| 19 | S | 134 | $0.12 \pm 0.01$ | $4.31 \pm 0.16$ | $6.63 \pm 0.20$ | $2.32 \pm 0.30$ | $5.61 \pm 0.71$ |
| 20 | T | 75 | $0.12 \pm 0.01$ | $4.35 \pm 0.30$ | $6.09 \pm 0.34$ | $1.73 \pm 0.37$ | $5.13 \pm 0.47$ |
| 21 | U | 13 | $0.07 \pm 0.01$ | $3.40 \pm 0.10$ | $5.43 \pm 0.13$ | $2.03 \pm 0.13$ | $3.64 \pm 0.05$ |
| 22 | V | 12 | $0.18 \pm 0.01$ | $3.24 \pm 0.47$ | $6.82 \pm 0.26$ | $0.42 \pm 3.57$ | $5.56 \pm 0.86$ |
| 23 | W | 14 | $0.12 \pm 0.01$ | $4.77 \pm 0.14$ | $6.57 \pm 0.10$ | $1.80 \pm 0.16$ | $5.50 \pm 0.64$ |

A total of 30 unique phrase types which were constructed by the iteration of 23 different note types were found in the song repertoire of Purple Sunbird (Table 2). Accumulation curves showed that the probability of finding new note is lesser compare to phrase (Figure 2 a and b).

**Figure 2.**
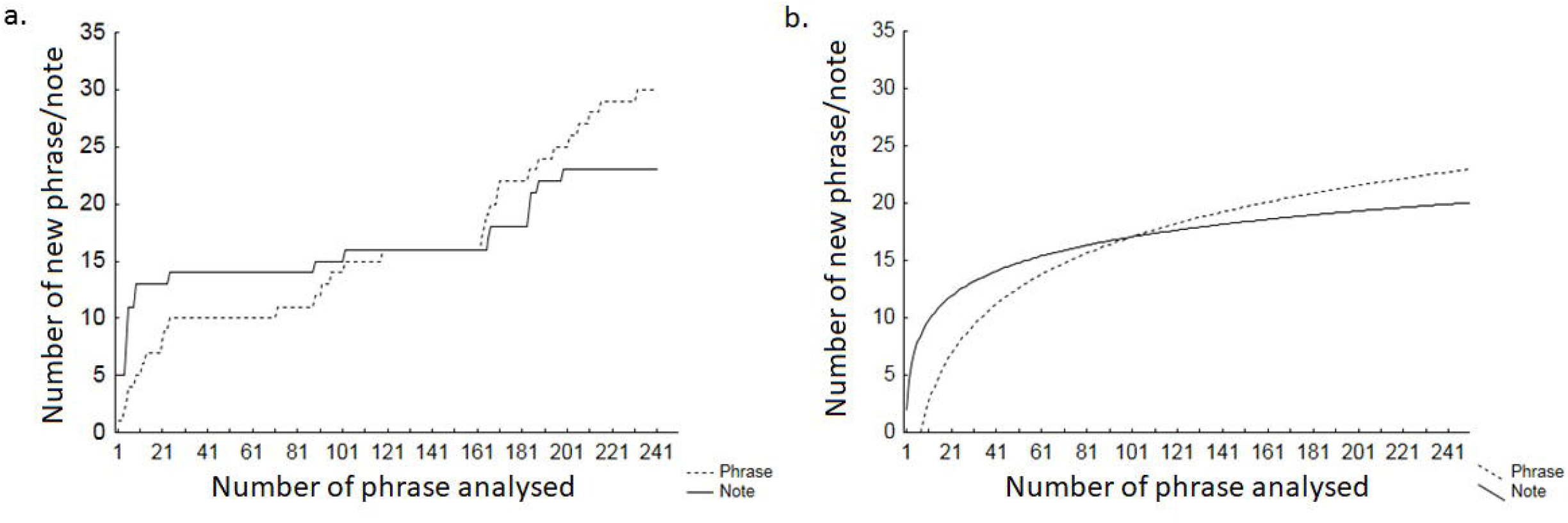
**a.** line plot and **b**. logarithmic curve showing accumulation of phrase and notes across 241 phrases sampled.

**Table 2.**
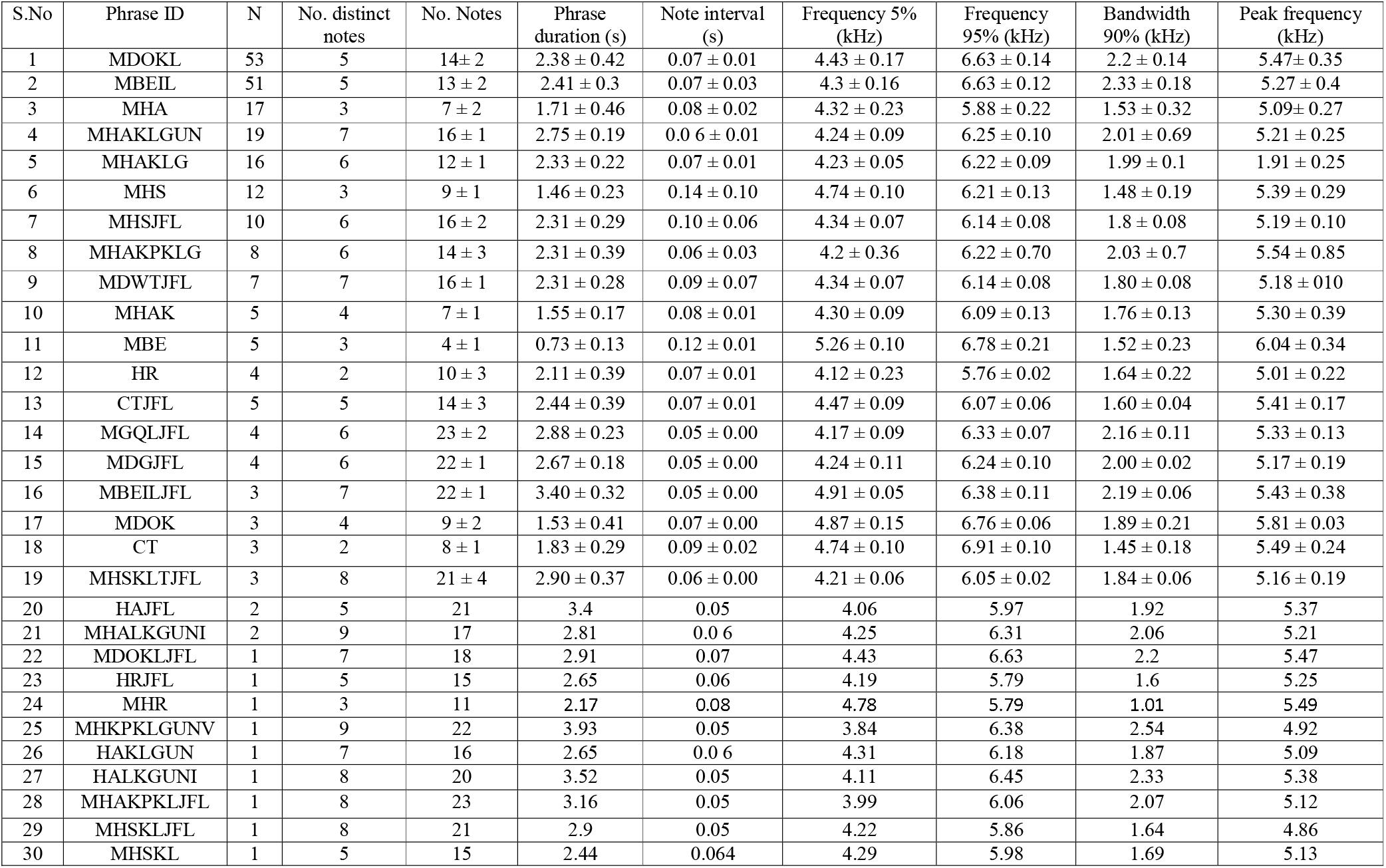
Mean ± SD of 7 acoustic parameters for 30 phrases obtained in the vocalization of Purple sunbird. N corresponds to sample size.

### Evidence of phonological syntax

The visual analysis found that each phrase initiated with a single introductory note or “prefix”, followed by series of notes consisting of 2-5 distinct note “phrase body” (Table 2). In some cases, there were additional notes (up to 3 acoustically distinct notes) at the terminal end of song phrase that we refer to as “suffix syllable” (Figure 3 a and b). We also found significant differences in percentage of occurrence of prefix versus suffix within phrases (χ^2^ = 95.12, df =1, p < 0.001) wherein prefix occurred in 93% of the phrases analysed whereas suffix syllable was found in only 27% of phrases (Figure 3 c). It was also found that various notes were restricted to a certain position (χ^2^ test; p < 0.001, Table S1) in the song phrase thereby reject the hypothesis that the notes occur at random order within the phrase. This implies that there is positional fidelity for notes. Prefix was restricted to note type “M”.

**Figure 3.**
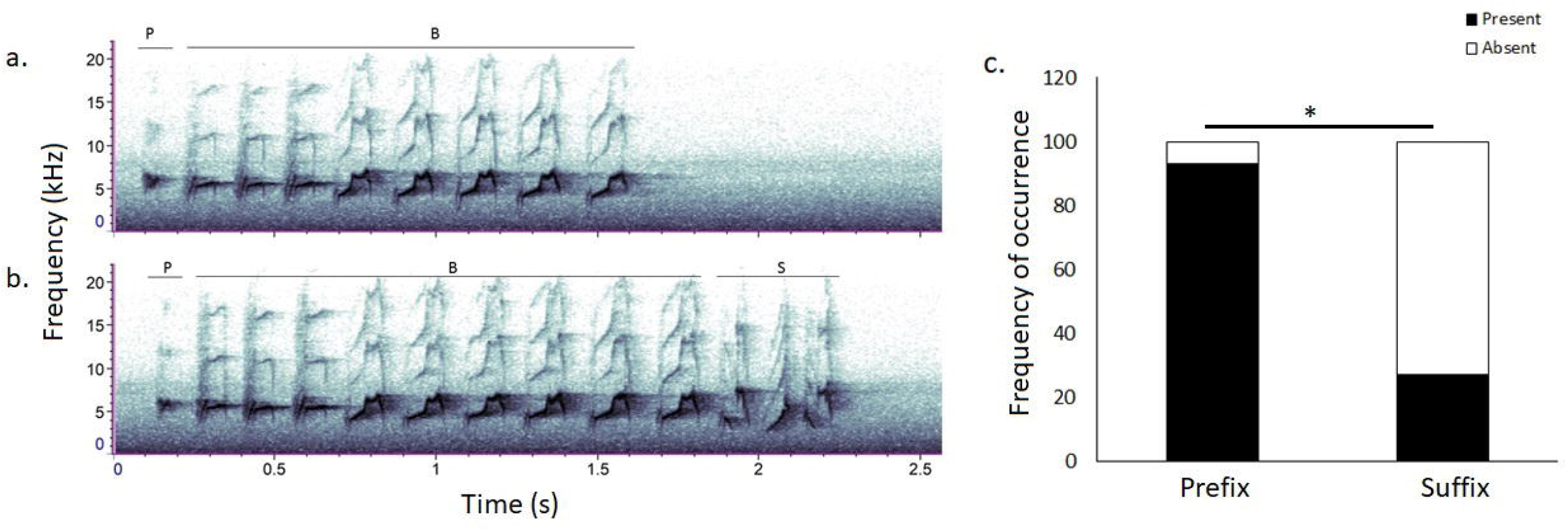
A representative song phrase of Purple sunbird; **a.** Phrase with prefix (P) and body(B); and **b.** Phrase with prefix (P), body (B) and suffix (S). **c**. Percentage of occurrence of prefix and suffix in song phrase. * represent significant difference.

Whereas, note types J and F and U and N always appear together respectively, forming the syllables JF and UN. Further, these two syllables were restricted to the suffix region of phrases. Similarly, note types C, B, D, H and T, R, G were restricted to the initial and terminal position of the phrase body respectively (Figure 4 a). The ‘expected’ heatmap shows that the percentage of occurrence of each note on a specific position is much higher than predicted by chance alone (Figure 4 b).

**Figure 4.**
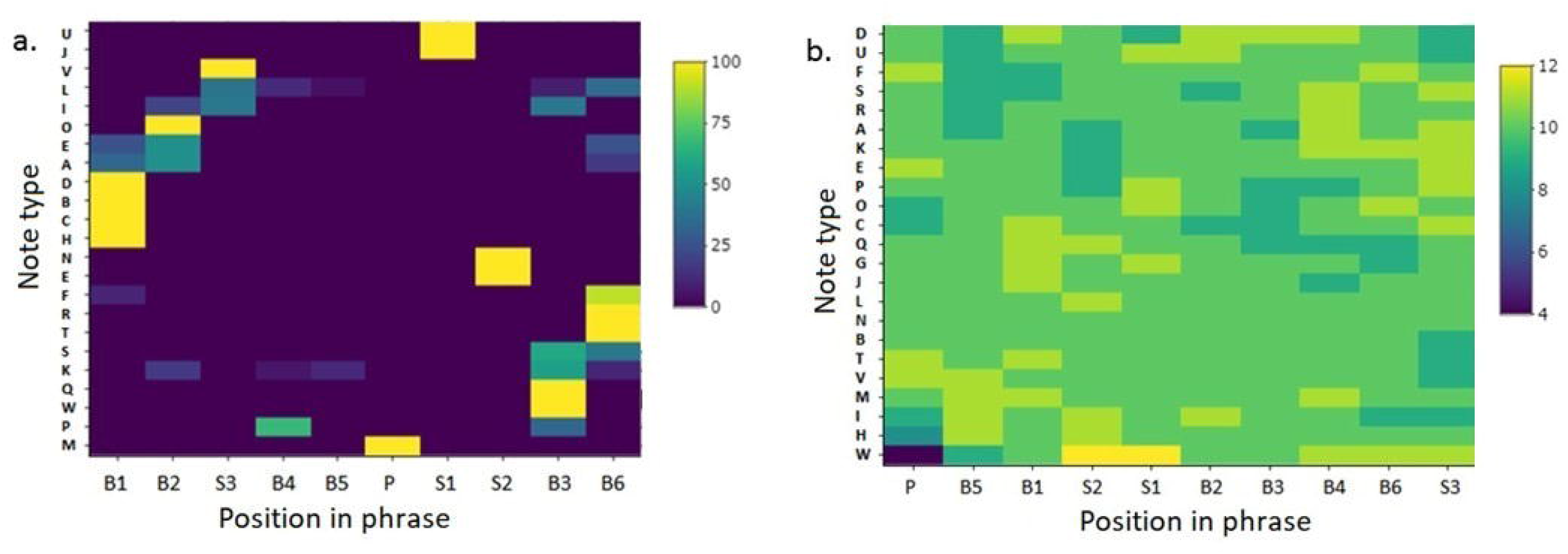
**a**. and **b.** Heat plots representing the percentage of occurrence of each note at specific positions within the phrase body: observed and expected respectively. Different color shades represent the percentage of occurrence of a note at a particular location in song phrases. P represent prefix, B represent body and S represent suffix. Numerical values along with B and S in x-axis represent position in ascending order,

### Evidence of complex vocal mechanism

The average inter-note duration between prefix and initial note of phrase body was 142 ± 50 (standard deviation (SD)) ms and within body was 63 ± 15 (SD) ms. The notes in suffix syllables were separated by a small inter-note time interval 14 ± 4 (SD) ms. We found that the there is a strong negative correlation between inter-note interval and its position (Pearson correlation: R = −0.54, t = −33.30, df =2652, p<0.001) in song phrase i.e., as the song phrase proceed, there is decrease in silent interval between the notes. This means that notes are delivered faster towards the end of a phrase. We also found there is significant difference in the temporal partitioning of notes with respect to its position within a phrase (GLM: p < 0.001). Significant difference was found in inter-note duration between PB and B (MW U test: W= 14, p < 0.001); PB and SS (MW U test: W= 300, p < 0.001); and between B and SS (MW U test: W= 415, p < 0.001) (Figure 5).

**Figure 5.**
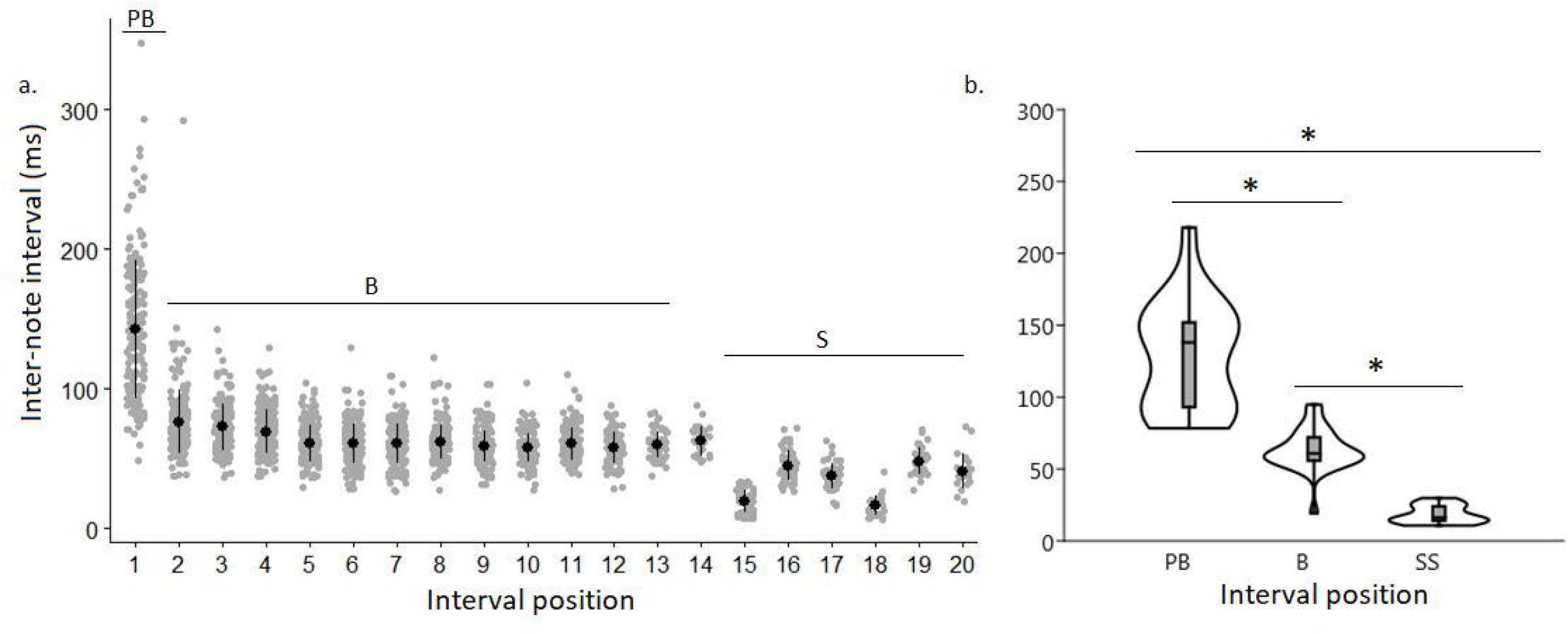
**a**. Distribution of inter-note interval within the song phrase of Purple Sunbird. Black dot represents mean and bar represents Standard Deviation. Grey dots show distribution of each data point. PB is inter-note interval between prefix and first note of phrase body, B is within body and S is within suffix inter-note interval. **b**. Violin plots representing differences in inter-note interval between prefix-body (PB), within body (B) and between suffix syllable (SS). The marker represents median, box represent interquartile range and spread is confident interval (0.95) in the plot. The shape of violin display distribution of each data point. * represent significant different.

## Discussion

The study provides the evidence for song complexity in terms of number of distinct notes and phrases in song repertoire, phonological syntax in construction of phrase and sexy-syllable in the breeding song of male Purple Sunbird. Based on the classical (aural-visual inspection) method, we found 23 structurally and aurally distinct notes which combine variably to form 30 different phrases. Number of unique notes based on classical classification vs CART analyses were 23 and 22 respectively. This implies that the results are in agreement with each other, thereby cross validating the two methods. Further, from accumulation curve, we found that the probability of finding new phrase is more compare to note. This is because phrases are constructed by iteration of existing notes and deletion and addition of note to the existing phrase result in addition of new phrase in the repertoire (Kroodsma 1977).

The presence of 30 different phrases in the breeding song repertoire of Purple Sunbird is relatively large when compared to other passerines 12 in Song sparrow (Hiebert et al. 1989), 12 in Chestnut-sided warbler (Byers 1995) and 4 in Great tit (McGregor et al. 1981). It is very likely that the size of song repertoire changes with individual as genetic, environmental, and cultural factors have great impact on the vocal performance of an individual (Nowicki et al. 2002; Reid et al. 2004; Roper and Zann 2006). Moreover, each song phrase of Purple Sunbird is found to be composed of 2-10 structurally and aurally distinct notes and the number of notes within a phrase varies from 3-24. According to Kroodsma 1977, song complexity depends upon number of distinct song component (note) within a song type (phrase) and in males Chaffinches complex song phrases with larger number of trills are selected by females (Leitão et al. 2006).

The song phrases of Purple Sunbird are composed of introductory note followed by series of notes arranged in stereotypic pattern (body) and later terminated by terminal trill (suffix). Moreover, each phrase type has fixed sequence of notes with occasional variation of addition or deletion of a note type. Thus, the syntax in Purple Sunbird song phrase is phonological than combinatorial, since the phrases themselves do not have a definite meaning and are composed of meaningless units, notes (Berwick et al. 2011). The entire song, composed of multiple repetition of phrases is used as a vocal display. Furthermore, the introductory note is a single note which is more comparable to that of single introductory whistle in the songs of male White-crowned Sparrows (Phuget sound) and Hermit Thrush (Roach et al. 2012; Nelson and Soha 2004) than to the series of introductory notes in the song of Zebra finches (Williams 2004). Introductory note in many species is a stereotypic note that occur at the onset of song or song phrase (Roach et al. 2012, Nelson and Soha 2004; Williams 2004) and is consistent in the song of Purple Sunbird as 93% of song phrase are initiated with an introductory note and was a stereotype note ‘M’.

We found that inter-note interval reduced from the start to end of a phrase. The prefix was observed to appear much earlier in the phrase, since inter-note time interval between prefix and first note of phrase body was 142 ms, whereas that of notes within phrase body was 63 ms. In suffix, within a 2-note syllable, each note (with different frequencies) was separated by a very small-time interval of 14 ms. Syllables comprising of multiple notes with such small inter-note interval, known as a mini-breath, are referred to as ‘sexy syllable’ (Garcia-Fernandez et al. 2013). It has been reported that females of the domestic Canary prefer males as mates which were able to incorporate two-note (each different frequency) syllables (separated by mini-breaths) into their songs (Garcia-Fernandez et.al 2013). Further, Suthers et al. (2004) reported that sexy syllables are hard to produce, which correlate with our findings as suffix syllables are present only in 27% of phrases. Thus, we speculate that suffix syllables (JF and UN) are also likely to be sexy syllables in Purple Sunbird. This, however, remains to be tested.

In conclusion, our findings revealed the presence of a large repertoire size in Purple Sunbird, in which phrase construction follows certain rules or phonological syntax. We also showed the presence of mini-breaths in song syllables. These results suggested that vocal communication in the Purple Sunbird is complex. Increase in complexity in vocalizations may also arise from variation in dialects owing to location or even seasonal variation. For instance, variation in songs based on individual and location has been reported in Song Sparrow (Harris and Lemon 1972), White-crown Sparrow (Nelson 2000) and Phuket Sound White-crown Sparrow (Nelson and Soha 2004). Whereas in free-ranging Canaries, song composition changes with season even though overall number of elements (notes/syllable) remain the same (Voigt et al. 2001). Studies also suggest that complexity in vocalizations is directly proportional to sexual attractiveness (Eriksson and Wallin 1986; Gil and Slater 2000) or is important in male-male competition (Ten Cate et al. 2002). Based on our findings, further studies can be carried-out in Purple Sunbird, as it is broadly distributed and is non-migratory in the tropics and subtropics region (Ali and Ripley 1983). Further, distinct sexual dimorphism during the breeding season also makes it an efficient system to study seasonal changes in the song repertoire.

## Acknowledgements

We thank IISER Mohali for infrastructural support. Both authors benefitted from several discussions during the poster session of IBAC 2017 where the initial results of the work were presented. Towards this we thank the organizers for arranging the conference in India, making it possible for many Indian students including the first author to attend it and benefit from discussion with an international audience.

## Ethical Statement

The study was carried out in the field and was purely observational. No animals were captured or harmed in anyway during the study.

