## Supplementary File for "Song complexity in relation to repertoire size and phonological syntax in the breeding song of Purple Sunbird"

| Note type | χ2 | df | p |
| --- | --- | --- | --- |
| A | 287.8 | 9 | <0.001 |
| B | 900 | 9 | <0.001 |
| C | 900 | 9 | <0.001 |
| D | 900 | 9 | <0.001 |
| E | 275 | 9 | <0.001 |
| F | 900 | 9 | <0.001 |
| G | 643 | 9 | <0.001 |
| H | 900 | 9 | <0.001 |
| I | 260 | 9 | <0.001 |
| J | 900 | 9 | <0.001 |
| K | 268.2 | 9 | <0.001 |
| L | 201.25 | 9 | <0.001 |
| M | 900 | 9 | <0.001 |
| N | 900 | 9 | <0.001 |
| O | 900 | 9 | <0.001 |
| P | 457.8 | 9 | <0.001 |
| Q | 900 | 9 | <0.001 |
| R | 900 | 9 | <0.001 |
| S | 420 | 9 | <0.001 |
| T | 900 | 9 | <0.001 |
| U | 900 | 9 | <0.001 |
| V | 900 | 9 | <0.001 |
| W | 900 | 9 | <0.001 |

Table S1. Results of Chi square test examining positional fidelity in notes within phrases.
